# Extensive cellular multitasking within *Bacillus subtilis* biofilms

**DOI:** 10.1101/2022.09.02.506450

**Authors:** Sarah M. Yannarell, Eric S. Beaudoin, Hunter S. Talley, Alexi A. Schoenborn, Galya Orr, Christopher R. Anderton, William B. Chrisler, Elizabeth A Shank

## Abstract

*Bacillus subtilis* is a soil-dwelling bacterium that can form biofilms, or communities of cells surrounded by a self-produced extracellular matrix. In biofilms, genetically identical cells often exhibit heterogeneous transcriptional phenotypes so that only subpopulations of cells carry out essential yet costly cellular processes that allow the entire community to thrive. Surprisingly, the extent of phenotypic heterogeneity and the relationships between subpopulations of cells within biofilms of even in well-studied bacterial systems like *B. subtilis* remains largely unknown. To determine relationships between these subpopulations of cells, we created 182 strains containing pairwise combinations of fluorescent transcriptional reporters for the expression state of 14 different genes associated with potential cellular subpopulations. We determined the spatial organization of the expression of these genes within biofilms using confocal microscopy, which revealed that many reporters localized to distinct areas of the biofilm, some of which were co-localized. We used flow cytometry to quantify reporter co-expression, which revealed that many cells ‘multi-task’, simultaneously expressing two reporters. These data indicate that prior models describing *B. subtilis* cells as differentiating into specific cell-types, each with a specific task or function, were oversimplified. Only a few subpopulations of cells, including surfactin and plipastatin producers, as well as sporulating and competent cells, appear to have distinct roles based on the set of genes examined here. These data will provide us with a framework with which to further study and make predictions about the roles of diverse cell phenotypes in *B. subtilis* biofilms.

**IMPORTANCE:** Many microbes differentiate, expressing diverse phenotypes to ensure their survival in various environments. However, studies on phenotypic differentiation have typically examined only a few phenotypes at one time, thus limiting our knowledge about the extent of differentiation and phenotypic overlap in the population. We investigated the spatial organization and gene expression relationships for genes important in *B. subtilis* biofilms. In doing so, we mapped spatial gene expression patterns and expanded the number of cell populations described in the *B. subtilis* literature. It is likely that other bacteria also display complex differentiation patterns within their biofilms. Studying the extent of cellular differentiation in other microbes may be important when designing therapies for disease-causing bacteria, where studying only a single phenotype may be masking underlying phenotypic differentiation relevant to infection outcomes.

## INTRODUCTION

Bacterial communities exist across diverse ecosystems. In these communities, genetically identical bacterial cells undergo differentiation that results in transcriptionally and functionally distinct cellular phenotypes (1). Such differentiation can result from nutrient availability (2), interspecies coculture interactions (3), stochastic effects (4), or specific microenvironments (5). It is thought that cells differentiate into phenotypically distinct subpopulations as a community survival strategy (2, 6–8). For many bacteria, the resulting phenotypic heterogeneity among genetically identical cells has implications for surface sensing (9, 10), virulence (11–13), and metabolism (14). For instance, some cell subpopulations may specialize in sugar incorporation (15) while others may release metabolic products (16). There is evidence of cross-feeding between glucose-fermenting and acetate-respiring subpopulations in *Escherichia coli* (17) and coordination between cells from the interior and periphery of *Bacillus subtilis* colony biofilms (18). Phenotypic differentiation is also implicated in antibiotic tolerance: *Pseudomonas aeruginosa* exhibits heterogeneity in its cellular metabolism due to oxygen gradients, which impacts how cells respond to antibiotics (19, 20). In *B. subtilis*, distinct subpopulations produce energetically costly compounds, like extracellular matrix components (21), extracellular proteases (22), or surfactin (23). This division of labor may have ecological benefits, since surfactin reduces surface tension to allow migration across solid surfaces (24) and extracellular matrix promotes plant root adherence (25). Given these potential incentives for cellular differentiation, phenotypic heterogeneity is a hallmark of bacterial biofilms, which are complex communities of cells encased by a self-produced extracellular matrix (26). Many models of cellular differentiation exist, but *Bacillus subtilis* is one of the best characterized, as well as being a genetically tractable, biofilm-forming bacterium (27). Based on the current (limited) fluorescent transcriptional reporter data available in the literature, *B. subtilis* is described as differentiating into six cellular phenotypes, leading to cells that are motile, matrix-producing, sporulating, cannibal, protease-producing, and competent (28) (Table 1). This model was derived based on data from fluorescent transcriptional reporters that use the expression of a marker gene as a proxy for the cell’s transcriptional state (e.g., a flagellar protein is a marker for motility), with heterogeneous gene expression leading to the designation of these cell types. Furthermore, the explicit examination of phenotypic overlap of these putative cell-types has been conducted in only a handful of cases (29–34). These studies indicate that motile, matrix-producing, and sporulating cells are spatiotemporally distinct within biofilms, as are matrix-producing cells from those that are competent or are producing the specialized metabolite surfactin. In contrast, matrix-producing and cannibal cells appear overlapping (23) and matrix-producing and protease-producing genes are co-expressed at certain times during growth (35). Beyond these few studies, the relationships between most other described cellular subpopulations in *B. subtilis* are unknown.

**Table 1.** *B. subtilis* phenotypes and corresponding genes

| Gene | Phenotype |
| --- | --- |
| *aprF* | Protease |
| *bacA* | Bacilysin |
| *comG* | Competence |
| *comQ* | Pheromone |
| *dhbA* | Bacillibactin |
| *hag* | Motility |
| *pksC* | Bacillaene |
| *ppsA* | Plipastatin |
| *sboA* | Subtilosin |
| *sdpA* | Cannibal (SDP) |
| *Skf* | Cannibal (SKF) |
| *skfAA* | Surfactin |
| *sspB* | Sporulation |
| *sunA* | Sublancin |
| *tapA* | Biofilm |

In addition to these six canonical cellular phenotypes, metabolite-producing cell subpopulations also play important roles in *B. subtilis* biofilm communities (36–38). *B. subtilis* NCIB3610 produces at least ten specialized metabolites, some of which act as cell-cell communication molecules and impact cellular differentiation (39). For instance, surfactin is a key factor in the formation of biofilm-matrix-producing cells (30, 39, 40), while ComX has been shown to stimulate the expression of surfactin through a ComP-ComA signaling cascade (41, 42). Of the remaining *B. subtilis* metabolites, four others have had intraspecific signaling bioactivities ascribed to them (23, 39, 40, 43, 44). While the expression of a handful of these metabolites have been previously examined for heterogeneous expression patterns in *B. subtilis* (23, 30, 45–50), only three of these have been examined within *B. subtilis* colony biofilms (23, 30, 50, 51), and even fewer have been examined in terms of their spatial organization (50). Thus, our understanding of the gene expression relationships between these specialized metabolites and the other described *B. subtilis* phenotypes is largely fragmentary, with many studies using cells grown under inconsistent growth conditions. Here we aim to examine the expression patterns of all ten metabolite biosynthetic genes and six currently described phenotypic cell-types under uniform biofilm-inducing conditions to obtain insights into the putative roles of these metabolites as cell-cell differentiation signals within *B. subtilis* model biofilms.

Considering the many described cell-types and metabolites known to exist within *B. subtilis* biofilms, the studies described above highlight that currently an incomplete understanding of *B. subtilis* cellular heterogeneity exists. We predict, based on the diverse gene expression relationships described so far, that substantial additional transcriptional multi-tasking occurs within *B. subtilis* biofilms. Here we aim to determine the extent of both transcriptional multi-tasking and the phenotypic cellular coordination within *B. subtilis* biofilms using single and pairwise fluorescent transcriptional reporters for genes previously associated with specific cell-types or specialized metabolites. Using flow cytometry and fluorescence microscopy, we quantitatively measured the expression overlap between the expression of these genes within individual cells as well as visualizing their spatial distributions within *B. subtilis* biofilms. Overall, we determined that some genes examined here were only expressed in a small subset of cells, while other cells multi-task, expressing multiple genes simultaneously. In addition, we observed that, overwhelmingly, most of the genes examined were expressed in distinct and repeatable spatial patterns across the biofilm. The data presented here thus provides a substantially improved, more comprehensive model of cellular heterogeneity within *B. subtilis* biofilms than currently exists.

## RESULTS

### Heterogeneous *B. subtilis* gene expression at the colony level

To monitor *B. subtilis* gene expression over time and space, we constructed strains containing fluorescent transcriptional reporters for key *B. subtilis* genes. To do this, we introduced the fluorescent protein Ypet (a variant of yellow fluorescent protein) under the control of promoters for 15 genes of interest (Table 1) and incorporated them into the neutral *amyE* locus (52). These genes are involved in specialized metabolite production, extracellular matrix production, motility, sporulation, competence, protease production, and cannibal antibiotic production. Exploring this wide range of genes was intended to provide a more global view of cellular phenotypic variation in *B. subtilis* than previously examined. We initially looked at the fluorescent transcriptional reporters for all ten of the specialized metabolites but excluded sublancin from future experiments because initial results showed that its colony morphology differed from that of wild type (Figure S1); we therefore focused on 14 *B. subtilis* genes.

To obtain information about the relationships between the expression of these genes, we first asked how the expression of these fluorescent transcriptional reporters were localized and how intensely they expressed *YPet* within *B. subtilis* biofilms. We grew biofilm colonies from an OD_600_-normalized inoculum on MSgg, a *B. subtilis* biofilm inducing media (53), and imaged colonies at 48 h (Figure 1) using brightfield and fluorescence illumination. We calculated the average fluorescence intensity across the colony, averaged from the center outward (Figure S2). At the colony level, we observed a range of reporter expression patterns. Two metabolite reporters (*sdpA* and *sboA*) were expressed at high levels; eight reporters (for genes encoding metabolite or structural products: *bacA, tapA, skfA, dhbA, comQX, hag, pksC*, and *sspB*) were expressed at mid-range levels; and the reporters for four genes (*ppsA, srfAA, comGA*, and *aprE)* were expressed at low or at near-background levels (Figure 1, Figure S2). With regards to their localization, the *sdpA, bacA, comQX, ppsA, srfAA*, and *aprE* reporters seemed to be consistently expressed throughout the colony (Figure 1, Figure S2). In contrast, *hag, sboA, dhbA*, and *pksC* were expressed primarily in the interior of the colony while t*apA*, *skfA*, and *sspB* were expressed mostly in the periphery (Figure 1, Figure S2). Thus, in comparing the fluorescence localization of different reporters at the colony level, we already observed some genes with similar spatial expression patterns and others with distinct expression patterns.

**Figure 1.**
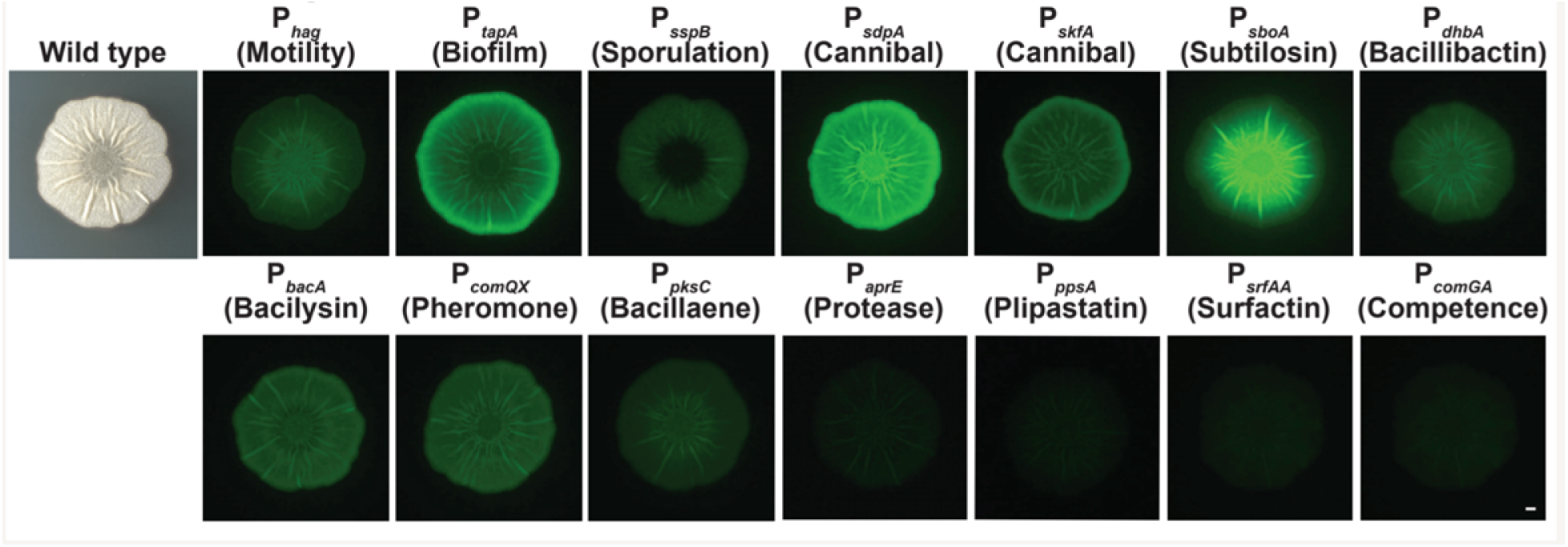
*B. subtilis* exhibits heterogeneous gene expression patterns at the colony level. Wild-type *B. subtilis* and *B. subtilis* strains containing fluorescent reporters were grown at 30 °C on the biofilm-inducing medium MSgg for 48 h. Brightness was linearly adjusted in the same way for each image using Fiji. Bar, 1 mm

### Identifying colony-level gene expression relationships using double-labeled strains

We hypothesized that the areas of the *B. subtilis* colony that appeared to be expressing more than one fluorescent transcriptional reporter could either be composed of highly heterogeneous populations of cells or else individual cells in those regions could be expressing multiple genes simultaneously, i.e., multitasking. To better visualize and quantify gene expression co-localization within *B. subtilis* colonies, we generated strains that contained pairwise combinations of these 14 reporters at two neutral sites on the *B. subtilis* chromosome (one reporter at the *amyE* locus (52) and the other at the *lacA* locus (54)) using phage transduction (Figure 2A). In combining reporter constructs for the 14 genes of interest, we created 182 strains; this included all 91 possible pairwise combinations of the 14 genes in each of the two color orientations (e.g., both P_*hag*_-*YPet*∷*amyE*, P_*tapA*_-*mTurq*∷*lacA* and P_*tapA*_-*YPet*∷*amyE*, P_*hag*_-*mTurq*∷*lacA* to control for differences in fluorescent protein expression levels). Only six such dual-reporter strain pairs (*hag-tapA*; *hag-sspB*; *tapA-sspB*; *srfAA-tapA*; *skfA-tapA*; and *tapA-comGA*) have been explicitly examined in the literature previously (23, 29, 30, 32, 33). All strains containing dual reporters grew similarly to wild-type *B. subtilis* (Figure S3). In analyzing the fluorescence levels of colony biofilms of these strains, we built a comprehensive picture of the spatial expression relationships between each of these gene pairs. We did so by growing the dual-reporter stains on MSgg and visualizing fluorescence within colonies at 48 h to directly compare expression patterns. In some pairs, like *sdpA* (cannibal) and *skfA* (cannibal), we observed high levels of co-localization (indicated by white in false-colored overlay image) (Figure 2B). Conversely, a *B. subtilis* strain containing *tapA* (matrix-producing) and *sboA* (subtilosin) reporters exhibited little co-localization between the expression of these two genes (Figure 2C).

**Figure 2.**
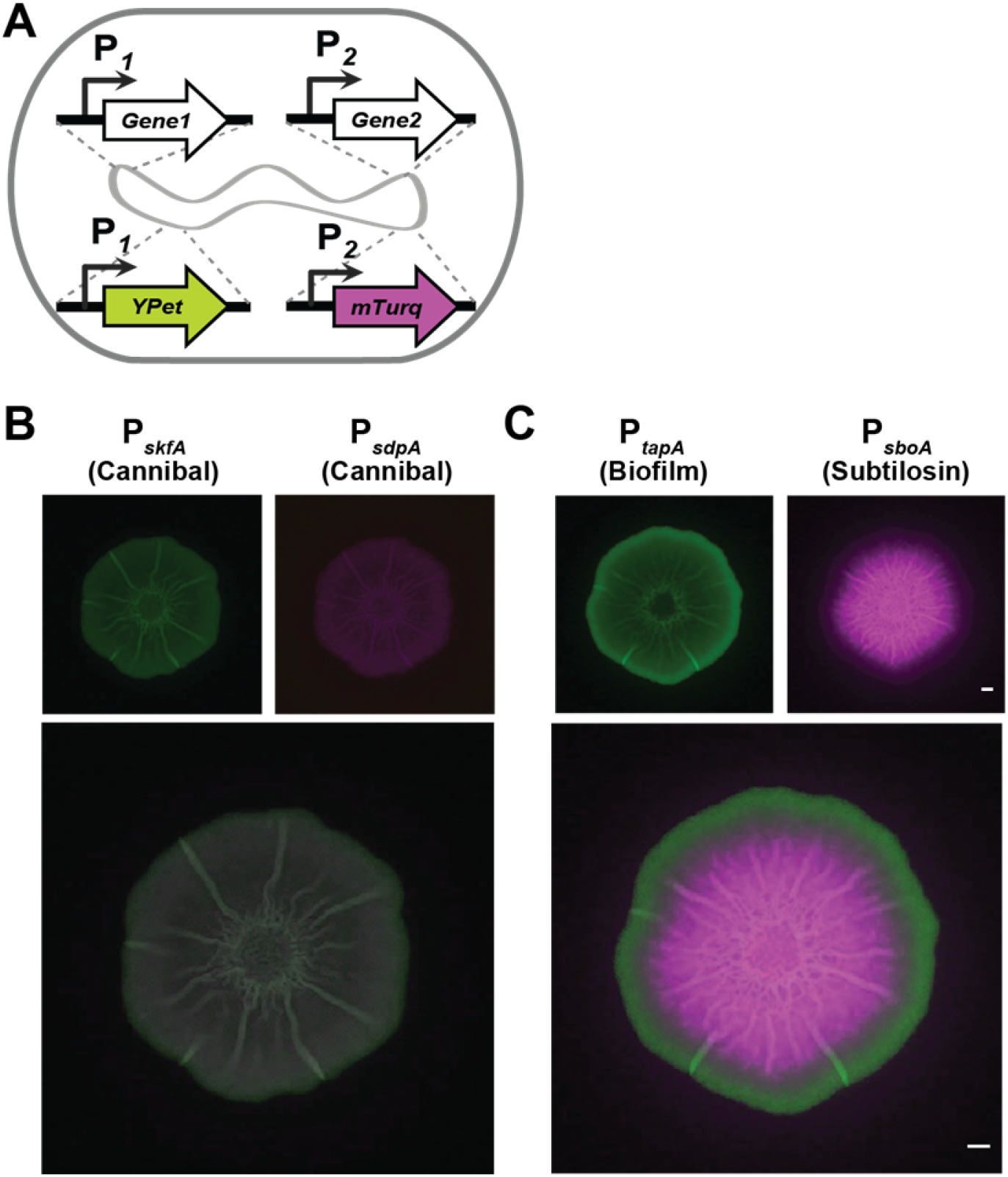
*B. subtilis* double reporters allow for direct localization comparison. A) Double reporter construction schematic B) Individual and merged channels of a *B. subtilis* biofilm containing *sdpA* (cannibal) and *skfA* (cannibal) reporters and C) a *B. subtilis* biofilm containing *tapA* (biofilm) and *sboA* (subtilosin) reporters grown on MSgg for 48 h. Colony images were taken from the top using a dissecting stereomicroscope. Bars, 1 mm.

### Analyzing gene expression in stratified, thin-sectioned colonies by confocal microscopy

The colony-level fluorescence microscopy allowed us to identify areas of the colony that appeared to contain co-localized reporter gene expression. We then delved further into this potential co-expression of genes using a confocal-laser-scanning microscope (CLSM) with an Airyscan detector. To gather spatial information not only from the surface of the colony, but also from individual cells within the biofilm, we quartered and thin-sectioned the colonies to 20 μm thick and flipped the sections on their side for imaging (schematic shown in Figure 3A). This approach enables a finer spatial visualization of fluorescence expression patterns within the depth of biofilms and provides information about the distributions of cells expressing different genes across the structure. For example, both *sdpA* (cannibal) and *skfA* (cannibal) reporters are present throughout the interior and peripheral regions of the colony and the reporters seem fairly well mixed in these regions, with some cells co-expressing both fluorescent reporters (white cells in Figure 3B). In contrast, *sboA* (subtilosin) and *tapA* (matrix-producing) are predominantly localized to the interior and the periphery, respectively, and these reporters seem to be mutually exclusive in individual cells (Figure 3C).

**Figure 3.**
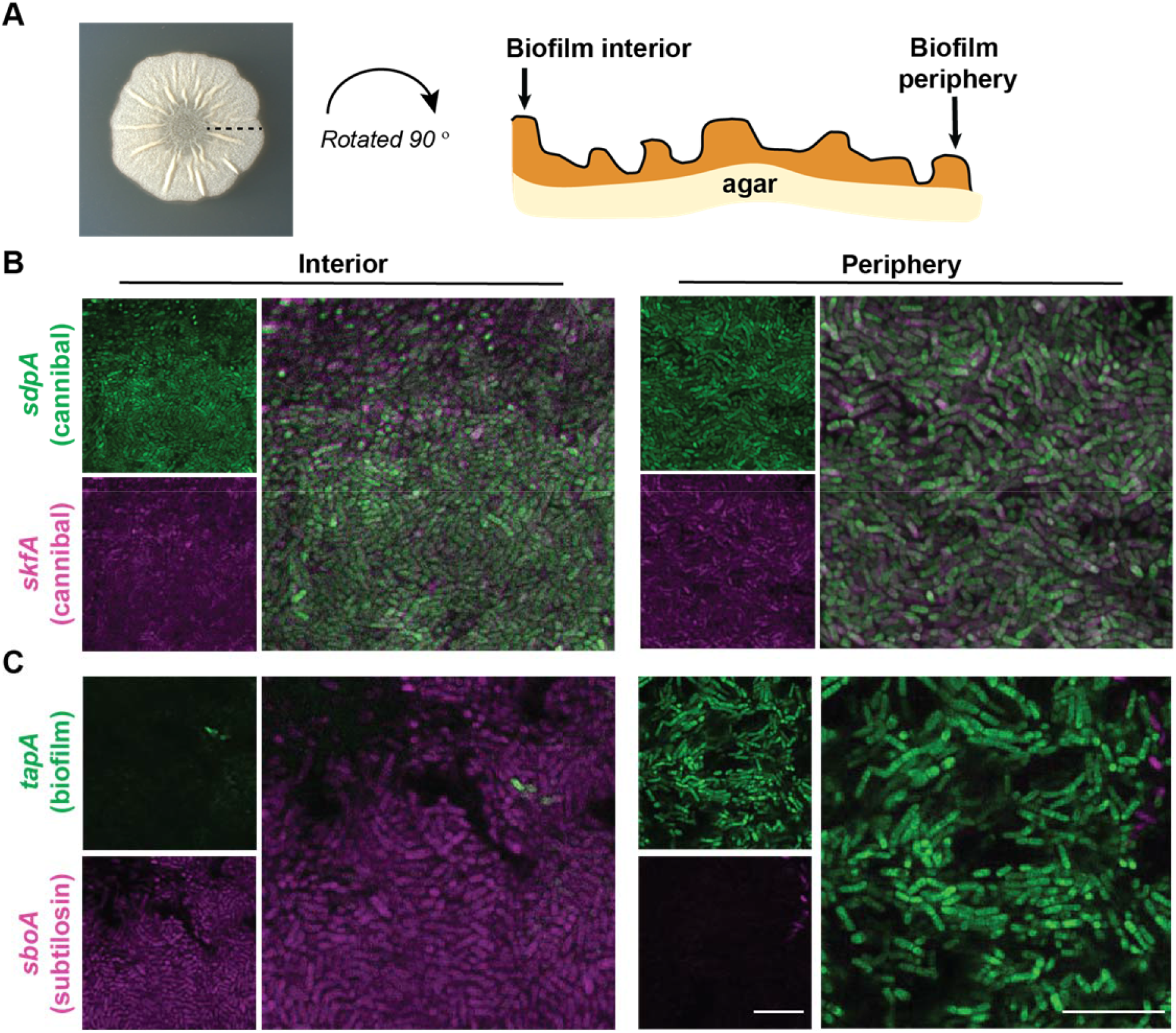
Specific phenotypic reporters display regions of co-localization and regions of distinct expression. A) Schematic of *B. subtilis* biofilm thin-section. B) Micrographs showing individual and merged channels of a *B. subtilis* biofilm containing *sdpA* (cannibal) and *skfA* (cannibal) reporters and C) *B. subtilis* biofilm containing *tapA* (biofilm) and *sboA* (subtilosin) grown on MSgg for 48 h, thin-sectioned, and imaged using Airyscan confocal microscopy at 100× to image interior and periphery regions. For each reporter pair, the intensities were optimized to show differences between reporters (i.e., intensities of B and C are not comparable). Bars, 10 um.

### Flow cytometry to quantify gene expression within individual cells

While the results from CLSM provided information about the spatial organization of gene expression across *B. subtilis* biofilm colonies, these data were not quantitative and in some cases it remained ambiguous whether (and the extent to which) genes were being co-expressed in the same cell. Therefore, we used flow cytometry to quantify gene expression and co-expression within *B. subtilis* biofilm cells using our double-labeled strains. We harvested colonies of all 182 dual-reporter strains grown on MSgg and fixed the cells using paraformaldehyde to prevent changes in expression levels during processing. Samples were sonicated, filtered, and analyzed on a flow cytometer, where data from a minimum of 24,000 cells per strain were collected.

We used *B. subtilis* non-fluorescent control samples to set the flow cytometry fluorescence detection gates, which enabled us to differentiate cell populations from each strain that were: a) not expressing either fluorophore, b) only expressing *mTurq*, c) only expressing *Ypet*, or d) expressing both fluorophores (Figure S4) To understand how many cells in the overall biofilm were expressing each individual gene, we first quantified the total percentage of cells expressing *Ypet* from each strain (Figure 4A). (We used only the *Ypet* signal from each dual-labeled strain for these calculations since the sensitivity of this fluorescent protein was superior to *mTurq* due to cellular background fluorescence in the *mTurq* channel). With this analysis, we determined that most cells within the *B. subtilis* population express *sdpA*, *comQX*, and *skfA*, while very few express *ppsA*, *srfAA*, or *comGA* (Figure 4A). The remaining eleven genes were expressed in between 25 - 75% of the cell population.

**Figure 4.**
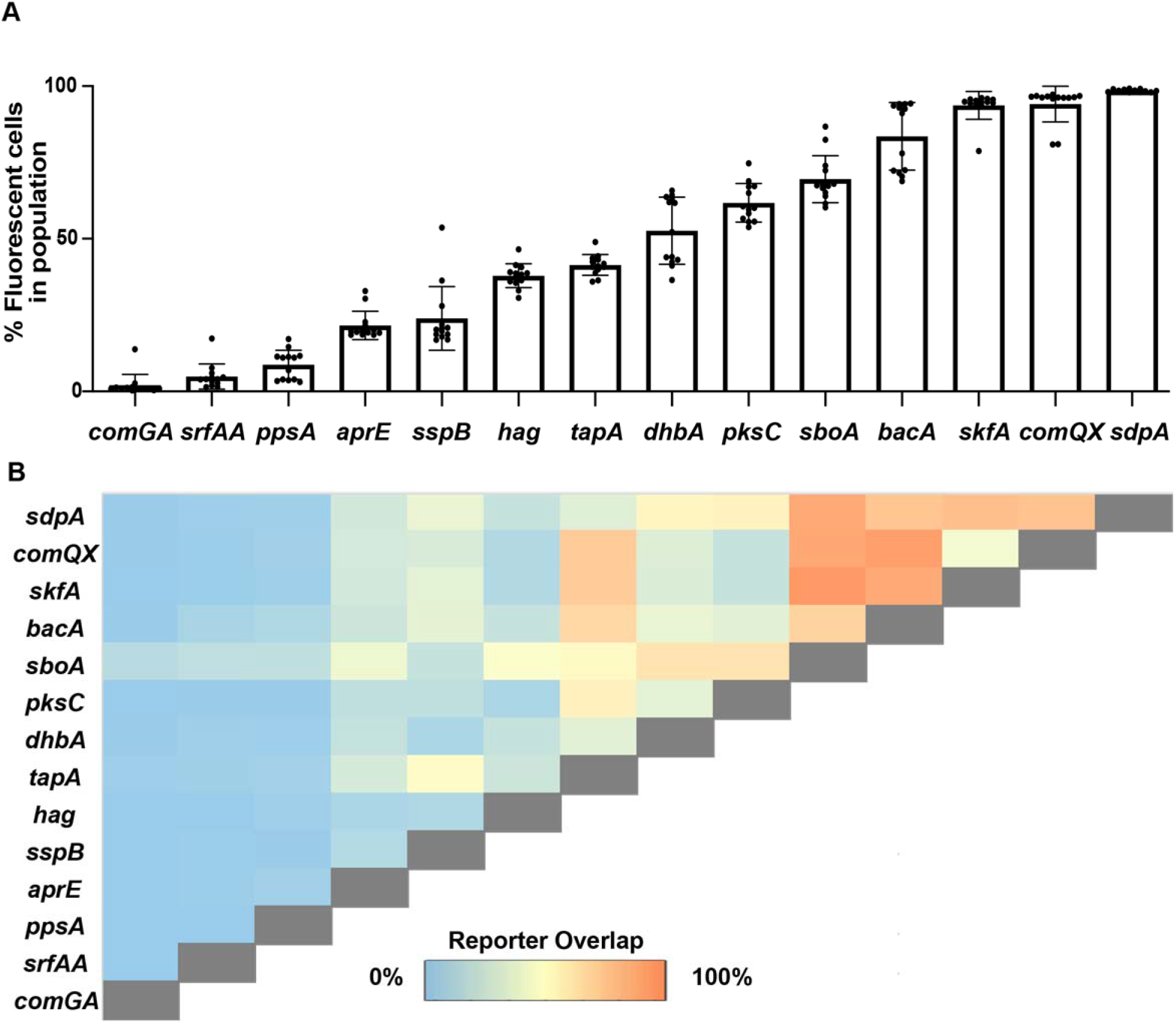
Many *B. subtilis* genes exhibit co-expression at the individual cell-level. A) Percent of fluorescent cells in a *B. subtilis* biofilm population after 48 h growth on MSgg was determined by flow cytometry. B) The percentage of cells co-expressing two reporters after 48 h growth on MSgg.

To understand the expression relationships between all of these genes, we then quantified the proportion of cells that expressed both *mTurq* and *Ypet* in each dual-labeled strain. We observed a range of gene-expression relationships, from completely distinct to fully overlapping (Figure 4B; all flow cytometry plots can be found in the Supplementary material (Figure S4)). Our dataset includes reporter pairs that corroborate results from previous studies (e.g., motility (*hag*) and sporulation (*sspB*) were not co-expressed (29) and biofilm matrix (*tapA*) and *skfA* (cannibal) reporters were overlapping (55)). However, in our data set, ~45% of the population co-express *tapA* and *sspB* reporters, which were originally described as distinct cell-types (29). Unexpectedly, *sdpA* did not appear to be coregulated with *tapA*, even though *tapA* is highly co-expressed with *skfA*, the other *B. subtilis* cannibalism toxin. Another unanticipated finding was that *comGA*, *ppsA*, and *srfAA* had minimal expression overlap with any other genes (Figure 4B) (with the exception of *sboA*, which was expressed in nearly all cells), indicating that cells expressing these genes may indeed represent more distinct cell-types that are specializing in a particular task. Beyond this, although most genes demonstrated some expression overlap with other genes (Figure 4B), some pairs of genes appeared to have anticorrelated expression: *sspB* and *dhbA, sspB* and *aprE, sspB* and hag, as well as *hag* and *aprE, hag* and *comQX*, *skfA*, and *pksC* (Figure 4B).

### Correlating the flow cytometry to microscopy

The flow cytometry results revealed extensive multitasking occurring in *B. subtilis* biofilm cells. However, by nature these flow cytometry data lack spatial information. We next wanted to investigate how cells expressing particular genes or gene pairs were spatially distributed within the colony. To do so, we used confocal microscopy to visualize representative reporter pairs with distinct flow cytometry distributions. In Figure 5 we show some examples of the diverse co-expression patterns we observed. For example, the flow cytometry data from the *sdpA*-*sboA* reporter pair indicate that almost all *sboA*-expressing cells also express *sdpA* (Figure 5A). In addition, however, there is a subset of *sdpA*-only expressing cells (Figure 5A, yellow bracket), and the cells expressing both reporters have a bimodal expression, where some express *sboA* at lower levels (Figure 5A, blue bracket), and others at very high levels (Figure 5A, red brackett). Interestingly, the spatial organization of these subpopulations of cells is not random: the brighter *sboA* population correlates with cells exhibiting higher expression at the colony agar interface (Figure 5D), while the *sdpA-*only subpopulation of cells are found at the colony-air interface (Figure 5D).

**Figure 5.**
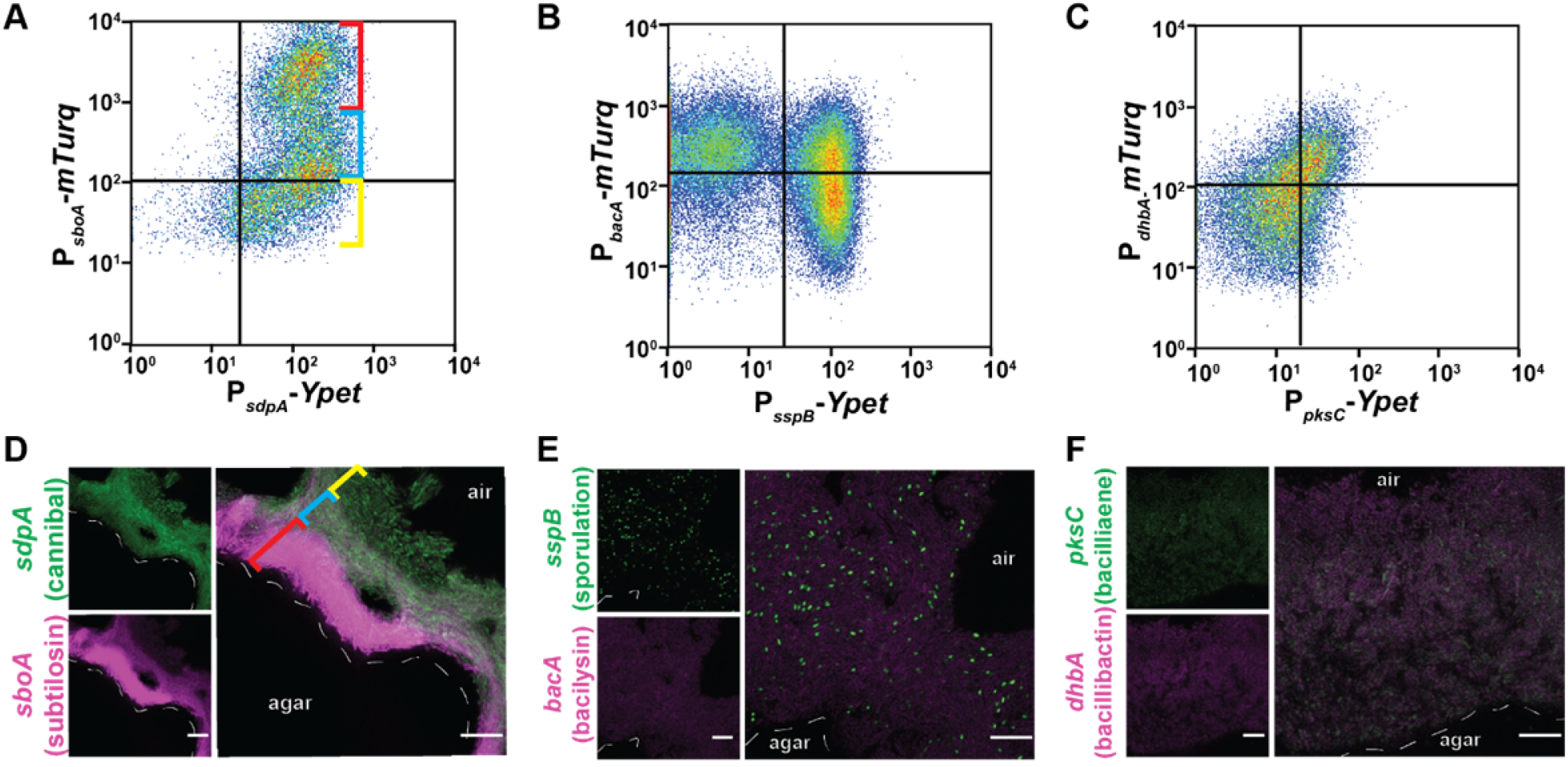
Corresponding reporter spatial arrangement with relationships displayed in flow cytometry data. Flow cytometry of the fluorescent intensities of *B. subtilis* cells containing A) *sdpA* and *sboA*, B) *sspB* and *bacA*, and C) *dhbA* and pksC reporters harvested from the 48 h timepoint. The gates were constructed from the non-fluorescent control sample from that experiment. A total of 24,000 cells were quantified for each sample. The *mTurq* reporter was detected using a 457 nm laser, and the *Ypet* reporter was detected using a 488 nm laser. Confocal microscopy of *B. subtilis* biofilm thin-sections containing D) *sdpA* and *sboA*, E) *sspB* and *bacA*, and F) *dhbA* and pksC reporters at 100×. For D and E, propyl gallate mounting medium was used. For each reporter pair, the intensities were optimized to show differences between reporters. Bars, 10um.

The *sspB* gene, a reporter for sporulation that is expressed during the first committed step of sporulation, exhibited an unusual, bimodal expression pattern. The *sspB* reporter was either on or off (Figure 5B). This is consistent with sporulation being known to be an all-or-none process at the stage at which the *sspB* gene is expressed; at 48 h of biofilm growth on MSgg we know that a subset of the population has begun to sporulate (29). The cells expressing *sspB* are also small and punctate in our fluorescence micrographs, indicative of cells undergoing sporulation (Figure 5E) (56, 57). In this field of view, we see little overlap with *bacA* (bacilysin)-expressing cells, which corresponds to the lower right quadrant of the flow cytometry data (Figure 5E); cells co-expressing *bacA* and *sspB* must reside elsewhere in the colony, since they are not visible here (Figure 5E).

Lastly, for a handful of reporters, we observed a partially overlapping gene expression pattern in our flow cytometry data (e.g., Figure 4B). The flow cytometry plot for reporter pair *dhbA* (bacillibactin) and *pksC* (bacillaene) displays this partial overlap, with each reporter expressed alone in a subset of cells as well as in some cells simultaneously (Figure 5C). Regions of co-localization in the center interior of the biofilm correlate to the observed co-expression in flow cytometry data (Figure 5F).

### Correlating reporters to each other using microscopy

We then examined the spatial relationships between different fluorescent reporters within each reporter strain of *B. subtilis* based on the confocal fluorescence micrographs. This was done as a voxel-based analysis using BiofilmQ (58). Voxels were 10 cubic pixels, or about 0.80 μm per side. Pearson correlation coefficients were derived for each voxel by looking at the intensity of each of the two reporters. Each Pearson value was overlaid onto the centroid location of the voxel. We saw that a range of distinct relationships were present (Figure 6) between different reporter pairs, supporting the diversity of relationships present between each set of genes as observed in the flow cytometry data. The expression of *sspB* and *sboA* were anti-correlated, while *pksC* and *bacA* expression was correlated in the center and on the edge of the biofilm. Interestingly, *pksC* and *dhbA* expression was correlated in the center but somewhat anticorrelated in the periphery, suggesting that a *pksC-* and *dhbA-*active cell-type is present in the center of the biofilm. These cross-sections provide an even more nuanced view of how different gene pairs are related to one another throughout the biofilm colony.

**Figure 6.**
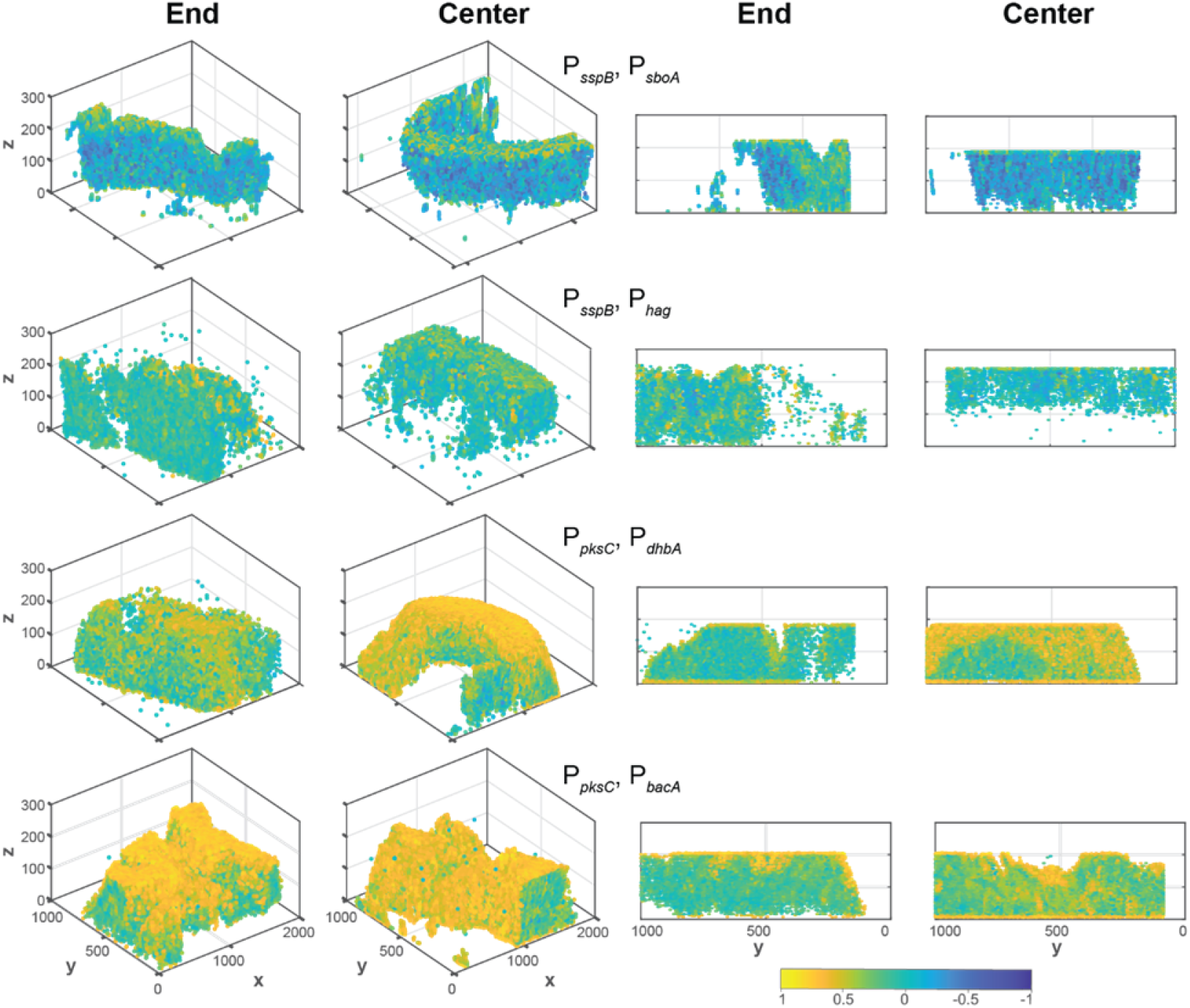
Correlations between pairs of reporters from confocal microscopy. Pearson correlation coefficients were determined between the fluorescent intensities of the two genes expressed in the dual-reporter strains of *B. subtilis*; the reporters are indicated for each row (ES338, ES329, ES392, and ES393). “End” indicates the peripheral, outer edge of the colony while “Center” indicates the center of the biofilm colony. The left two panels are three-dimensional representations of the colony slice and the right two panels are cross-sections of these images. Yellow indicates highly correlated fluorescence between the two reporters within each strain, while dark blue indicates anti-correlated fluorescence. Areas with no fluorescence are not represented in these images.

## DISCUSSION

Historically, researchers have classified *B. subtilis* into several cell states or subpopulations that were identified based on the gene expression inferred by a handful of fluorescent transcriptional reporters (36). In this study, we have built upon these foundational studies and generated 182 strains containing all pairwise combinations (91 reporter pairs in both color combinations) of a suite of 14 fluorescent transcriptional reporters that report on the gene expression of described cell physiologies as well as genes that encode specialized metabolite machinery in *B. subtilis*. By analyzing the spatial and co-expression relationships of genes controlling these critical phenotypes and the production of cell-cell metabolite signals within *B. subtilis* biofilms, we have uncovered which genes are simultaneously expressed within *B. subtilis* cells, enhancing our understanding of how extensively cells within these biofilm communities are multitasking. Furthermore, the expression patterns of these genes are spatially distributed in a repeatable way across biofilms (shown here and in (50)). Our ambitious evaluation of these many strains by both flow cytometry and microscopy has led to a substantially more nuanced and thorough view of cellular heterogeneity within *B. subtilis* biofilms.

Our study reveals a substantial level of multitasking within *B. subtilis* biofilm cells. We frequently observed subpopulations of cells that co-expressed two reporters as well as subpopulations that only expressed one of the two reporters (Figure 4). Our data indicate that 66% (60/91) of the phenotypes examined here are co-expressed in at least some *B. subtilis* biofilm cells (Figure 4F). Given the diverse reasons cells differentiate (e.g., division of labor in producing extracellular matrix or in generating cells tolerant to antibiotics, etc.), it is possible that *B. subtilis* cells may multitask to provide an additional layer of ‘bet-hedging’ in the face of environmental stress. Further work interrogating the broader, genome-wide gene expression patterns of individual cells within biofilms would be informative here. The global transcriptome of *B. subtilis* has been analyzed previously (59, 60), but data about the specific transcriptional profile of multitasking cells are masked when biofilms are harvested and analyzed in bulk. Previous studies have coupled fluorescence-activated cell sorting followed by RNA sequencing to examine competent (*comG*+) and non-competent subpopulations (61) and succinate co-A ligase (*sucC*+) populations (16) in *B. subtilis*. We anticipate using such approaches, in conjunction with our suite of dual-labeled fluorescent reporter strains, to isolate and determine the transcriptome of specific subpopulations of *B. subtilis* cells. Expanding the current collection of reporters to include other genes implicated in cell-cell communication and biofilm formation (62–65) would provide an even more refined view of cellular transcriptional heterogeneity within bacterial biofilms.

Similar to how the spatial arrangements of bacteria in multispecies biofilms impact the function and bioactivities of these communities (66–72), we expect that in single-species biofilms the spatial organization of cellular subpopulations (as defined by their distinct gene expression profiles) may also be important for the function of differentiated cells. The organization of cells with specific gene expression patterns may also provide clues about the underlying regulation of these genes. We expect that genes expressed in distinct locations across the biofilm may indicate that their expression is regulated by microenvironmental factors. For instance, the organization of *sdpA* and *sboA* reporters (Figure 4B) may indicate that *sdpA* expression is important for all cells, while *sboA* expression is more important near the agar; conversely, it may be important that *sboA is* not expressed at the top of the biofilm. In contrast, when two reporters are expressed in the same location but in different cells, for instance *pksC* and *dhbA* (Figure 4D), we predict that their regulation may not be microenvironment-specific but could instead be stochastic or directed by a positive feedback loop (73). A technique developed to use stochastic pulsing force the expression of genes typically not co-expressed (74) could be applied to investigate the underlying regulation of co-localized reporters in biofilms and determine the functional effects of dysregulated gene expression. Mutations could similarly be used to dysregulate gene expression; mutants in motility-related genes (*hag, cheA, cheY*) have been seen to alter the expression of *hag* (using a P_*hag*_-*yfp* reporter) (29).

*B. subtilis* and many other bacteria dedicate a large portion of their genome to specialized metabolite gene clusters (75, 76); many of the metabolites produced by these biosynthetic genes act as cell-cell signals (77). Many of the biosynthesis genes for specialized metabolites examined here were co-expressed and co-localized with other specialized metabolite and physiological reporter genes. This overlap suggests that the gene expression of the biosynthesis machinery for generating specialized metabolites such as subtilosin, bacillaene, bacillibactin, and comX are co-expressed with (and potentially regulate the generation of) cellular phenotypes such as matrix-production, cannibalism, and extracellular protease production. Other genes that exhibited minimal gene expression overlap (such as those for motility, sporulation, competence, plipastatin, and surfactin) may have more specific roles in the biofilm. Our work greatly advances our understanding of which cells within a biofilm express specialized metabolite biosynthesis genes, but only by using techniques that directly detect metabolites, such as our recent study using mass spectrometry imaging (50), are able to resolve how far these metabolites signals physically diffuse from the producing cells. Modeling approaches that are able to integrate all of these diverse datasets (microscopy, quantitative flow cytometry, mass spectrometry imaging) may enable us to generate a comprehensive and predictive spatial model of specialized metabolite signaling in bacterial biofilms.

This work greatly advances our understanding of the heterogeneity of cellular gene expression and transcriptional multitasking that exists within the biofilms of the model bacterium *B. subtilis*. The spatial transcriptional complexity we describe here within this single-species biofilm is likely to be further modified by interactions with other bacteria and fungi. We know that multiple other bacteria can affect *B. subtilis* physiology based on single-gene reporter constructs (44, 78, 79) and predict that these changes are representative of shifts in the balance of transcriptional heterogeneity that are propagated across many other genes. The tools and approaches we implemented here could similarly be utilized to address the question of cellular transcriptional heterogeneity within multispecies communities and how their spatial organization may shift in response to environmental stressors. This research, which provides an unusually complete depiction of how *B. subtilis* differentiates within biofilms, provides a foundation for exploring more complex metabolic and regulatory interactions between cells within environmentally and agriculturally important microbial communities.

## ACKNOWLEDGEMENTS

We gratefully acknowledge the assistance and advice of Srigokul (Gokul) Upadhyayula (UCBerkeley) on the image processing, troubleshooting, and computational analysis performed in this paper. This research was supported by funds provided by the National Institutes of Health (GM112981 to E.A.S.) A portion of this research was performed on a project award (10.46936/expl.proj.2019.51105/60000139) from the Environmental Molecular Sciences Laboratory, a DOE Office of Science User Facility sponsored by the Biological and Environmental Research program under Contract No. DE-AC05-76RL01830. Some of this material is based upon work supported by the U.S. Department of Energy, Office of Science, Office of Workforce Development for Teachers and Scientists, Office of Science Graduate Student Research (SCGSR) program (to S.M.Y). The SCGSR program is administered by the Oak Ridge Institute for Science and Education (ORISE) for the DOE. ORISE is managed by ORAU under contract number DE-SC0014664. All opinions expressed in this paper are the author’s and do not necessarily reflect the policies and views of NIH, DOE, ORAU, or ORISE.

## MATERIALS AND METHODS

### Bacterial strains and growth conditions

Table S1 lists the strains used in this study. *B. subtilis* strains were cultured on lysogeny broth (LB)-Lennox medium (10 g/liter tryptone, 5 g/liter yeast extract, 5 g/liter NaCl, 1.5% agar) at 30 °C for 16 to 18 h with antibiotics as necessary. TY broth consisted of LB supplemented with 10 mM MgSO_4_ and 100 μM MnSO_4_ after autoclaving. Colony biofilms were grown on MSgg medium (5 mM potassium phosphate [pH 7], 100 mM morpholinepropanesulfonic acid [MOPS; pH 7], 2 mM MgCl_2_, 700 μM CaCl_2_, 50 μM MnCl_2_, 50 μM FeCl_3_, 1 μM ZnCl_2_, 2 μM thiamine, 0.5% glycerol, 0.5% glutamate) at with 1.5% agar at 30 °C for 48 h. Antibiotics (final concentrations) were as follows unless noted otherwise: MLS (1 μg/ml erythromycin, 25 μg/ml lincomycin) and chloramphenicol (5 μg/ml). *Escherichia coli* strains were cultured in LB-Miller medium (10 g/liter tryptone, 5 g/liter yeast extract, 10 g/liter NaCl, 1.5% agar). Final concentration of carbenicillin was 50 μg/ml.

### Colony morphology phase contrast and fluorescence imaging

Macrocolony biofilm images were gathered using a Zeiss SteREO Discovery.V8 dissecting stereomicroscope (Zeiss, Oberkochen, Germany) with a 1 × 0.63 lens objective in three channels: brightfield, Ypet, and mTurq. Images were exported to as a .TIFF for image analysis at 1388 × 1040 pixels. These images were indexed and grouped based on the strain and reporters the strain contained. Images were then imported into Matlab 2020b. For each strain, the location of the colony was determined by masking the image using the Laplassian of Gausian of a grayscale brightfield image. This method of detection was used because the agar was a single, relatively uniform value, while the colony was a substantially different, relatively uniform value. The center of the colony was determined using the Centroid function in Matlab. The edge of the colony was determined using bwboundries function in Matlab on the masked colony image. The average Euclidian distance, in number of pixels, between the center and edge of the colony was then determined. Each pixel’s “Euclidian distance from the center of the colony” was divided by the “average Euclidian distance from the center of the colony to the edge,” transforming each pixel’s distance into a percentage (with the edge of the colony being 100%) to allow for easier comparison between colony images. Each distance was rounded to two decimal places, then the mean signal in mTurq and Ypet was taken for each unique distance at and graphed. Graphs were limited to 120% on the X-value to capture background fluorescence just beyond the edge of the colony. Y-values were normalized to an appropriate and consistent value for each reporter. The code can be found at DOI: 10.5281/zenodo.4624987.

### SPP1 phage transduction

Phage transduction was carried out as previously described (80). Briefly, we grew the *B. subtilis* donor strain at 37 °C in TY broth until the culture reached an OD_600_ of 1.0. At that point, we infected cells with SPP1 phage stock and incubated for 15 min at 37 °C. We then added 0.5% TY soft top agar to the cells and phage, overlaid the mixture on TY 1.5% agar plates, and incubated plates at 37 °C for 8 to 16 h. *B. subtilis* donor phage plaques were collected and pelleted using a clinical centrifuge. We infected *B. subtilis* recipient cells with three hundred microliters of supernatant, and then plated the cell lysate on LB-Lennox with 10 mM citrate and antibiotic to which the donor strain was resistant. Plates were incubated at 37 °C for 12 to 24 h. Four colonies were picked from each phage transduction and struck on LB-Lennox plates with antibiotic. After growth, strains were restruck two more times on LB-Lennox plates with antibiotic. Cells were spotted on MSgg and incubated at 30 °C to ensure growth, which indicates the cells have a 3610 rather than a 168 background (which is an amino acid auxotroph). Specifics of reporter construction are described below.

### Construction of *B. subtilis* reporter strains

The newly constructed transcriptional reporter plasmids (pES099 – pES112) containing *Ypet* were derived from pES045 (*amyE:*:P_*spacC*_-*Ypet*) (81, 82). To construct these plasmids, the *spacC* promoter was removed by digestion with EcoRI and HindIII and replaced with promoter sequences. Promoter sequences were amplified from *B. subtilis* wild-type genomic DNA (See primers in Table S1) and inserted into the base plasmid by isothermal assembly (also used for all subsequent constructions described in this section) (83) and transformed into *E. coli*.

A plasmid containing mTurquoise2 (*mTurq*) was generated using primer ES395 and primer ES315 (see Table S1) to amplify *mTurq* from GL-FP-31. The fragment was cloned into plasmid pDR183 [*lacA*∷(*mls*)] (84) digested with SalI and EcoRI. To create *mTurq* reporters, we amplified promoter sequences from *B. subtilis* wild-type genomic DNA (see primers in Table S1), digested with NheI and SalI, and inserted into the pDR183-*mTurq* base plasmid (pES069) using Isothermal assembly. The assembled plasmids were transformed into *E. coli*.

Upon final construction, the plasmids were isolated from *E. coli*, linearized, and transformed into *B. subtilis* 168 cells grown to stationary phase. Cells containing *Ypet* reporters were plated on Lennox-chloramphenicol to select for transformants. Cells containing *mTurq* reporters were plated on Lennox-MLS to select for transformants. Phage transduction was carried out as described previously (80) and above. *B. subtilis mTurq* reporters were used as the donor strains and grown to 37 °C in TY broth until the culture reached an OD_600_ of 1.0. Cells were infected with SPP1 phage stock and plated on 0.5% TY soft top agar, overlaid on TY 1.5% agar plates, and incubated at 37 °C for 8 to 16 h. *B. subtilis* donor phage plaques were collected and pelleted using a clinical centrifuge. Three hundred microliters of supernatant was used to infect *B. subtilis* 3610 wild-type and *B. subtilis Ypet* reporter strains (recipient cells) to construct single and dual-fluorescent reporters, respectively. The cell lysate was then plated on LB-Lennox with 10 mM citrate and MLS to which the donor *mTurq* reporter strains were resistant. Plates were incubated at 37 °C for 12 to 24 h. Three colonies were picked from each phage transduction and struck on LB-Lennox plates with MLS and citrate to select for *B. subtilis* cells that contained *mTurq* reporters. For strains containing dual-fluorescent reporters, strains were then re-struck on Lennox-chloramphenicol to select for strains containing both *mTurq* and *Ypet* reporters. Cells were spotted on MSgg and incubated at 30 °C to ensure growth, which indicates the cells have a 3610 rather than a 168 background (which is an amino acid auxotroph). Colony morphology of reporter strains were compared to wild type, as morphology should be identical.

### Flow cytometry

*B. subtilis* strains were prepared and grown on MSgg as described above. After 48 h of growth, biofilms were collected and resuspended in 1 mL 1× phosphate buffered saline (PBS) using a 23G needle and syringe to shear the biofilm. Cells were pelleted by centrifugation at 16,000 × g, the supernatant was removed, and the cells were fixed in 200 μL of 4% (w/v) paraformaldehyde for 7 minutes. After the incubation, the cells were pelleted, washed in 1× PBS to remove residual paraformaldehyde, and resuspended in GTE buffer [1% glucose (wt/vol) and 5 mM EDTA in 1× phosphate buffer, pH 7.4]. Samples were stored at 4 °C until flow cytometry analysis. Prior to analysis, cells were sonicated for 12 pulses [1 s pulse with subsequent 1 s pause] and filtered through a 38-μm nylon mesh. *Ypet* and *mTurq* fluorescence in dual-reporter strains was measured using the 488 and 457 lasers, respectively, of the Influx cell sorter (BD Biosciences).

### Thin-sectioning

The thin-sectioning protocol was adapted from Vlamakis et al. and Marlow et al (29, 35). *B. subtilis* strains were cultured on MSgg as described above and vapor fixed with 8% paraformaldehyde (adapted from (85)). Biofilm-agar blocks were quartered, transferred to a 15 × 15 × 5 mm mold (Fisher; Cat: 22-363-553), and snap-frozen. The colony was then overlaid with 4% (w/v) agarose (Lonza; Cat: 50181) and frozen at −80 °C for 15 min. The blocks were transferred to −20 °C for 30 min to equilibrate. Colony blocks were mounted to the chuck with double-distilled H_2_O and sliced to 20 μm thick cross-sections using a cryotome (Thermo cryostar NX70). Sections were attached to VWR Superfrost Plus slides (Cat: 48311-703), and stored at −20 °C.

### Confocal microscopy and image analysis

Sections for microscopy were overlaid with mounting medium ProLong Gold Antifade Mountant (ThermoFisher; Cat: P10144) unless otherwise noted and a 25 × 25 mm coverslip (Fisher; 12-548-C). Sections were imaged using a Zeiss 710 laser scanning confocal microscope equipped with a 20× EC Plan NEOFLUAR and 100× Plan APOCHROMAT oil objective.

## DATA AVAILABILITY

The code for spatial quantification of fluorescence over the bacterial colonies can be found at DOI: 10.5281/zenodo.4624987. Images of all flow cytometry plots are available in the supplemental material section as Figure S4. All raw microscopy images will be uploaded onto Dryad.com (currently in process).

## SUPPLEMENTAL MATERIAL

**Table S1**.

Table S1 lists the strains used in this study.

**Figure S1**. Colony morphology of *B. subtilis* wild-type and *B. subtilis* P_*sunA*_-*Ypet* grown at 30°C on the biofilm-inducing medium MSgg for 48 h. Colony images were taken from the top using a dissecting stereomicroscope. Bar, 1 mm.

**Figure S2**. Average fluorescent pixel intensity of *B. subtilis* biofilms containing single *Ypet* reporters depicted in Figure 1. Intensity is plotted (black line) from the center (x-axis = 0) to the edge of the colony (x-axis = 100). The baseline autofluorescence detected in the agar is displayed by the dashed red line. 95% confidence intervals are indicated by grey dashed lines.

**Figure S3**. Growth of wild-type and a representative subset of single and double reporter *B. subtilis* strains measured by OD_600_ measurements over time. Baseline corrected using blank MSgg medium.

**Figure S4**. Flow cytometry profiles of *B. subtilis* reporters pairs.

